# ALPACA: a fast and accurate approach for automated landmarking of three-dimensional biological structures

**DOI:** 10.1101/2020.09.18.303891

**Authors:** Arthur Porto, Sara M. Rolfe, A. Murat Maga

**Affiliations:** Center for Development Biology and Regenerative Medicine, Seattle Children’s Research Institute, Seattle, Washington; Friday Harbor Laboratories, University of Washington, San Juan Island, Washington; Division of Craniofacial Medicine, Department of Pediatrics, University of Washington, Seattle, Washington

**Keywords:** automation, phenotyping, phenomics, biological structures, image registration, open-source

## Abstract

Landmark-based geometric morphometrics has emerged as an essential discipline for the quantitative analysis of size and shape in ecology and evolution. With the ever-increasing density of digitized landmarks, the possible development of a fully automated method of landmark placement has attracted considerable attention. Despite the recent progress in image registration techniques, which could provide a pathway to automation, three-dimensional morphometric data is still mainly gathered by trained experts. For the most part, the large infrastructure requirements necessary to perform image-based registration, together with its system-specificity and its overall speed have prevented wide dissemination.

Here, we propose and implement a general and lightweight point cloud-based approach to automatically collect highdimensional landmark data in 3D surfaces (Automated Landmarking through Point cloud Alignment and Correspondence Analysis). Our framework possesses several advantages compared with image-based approaches. First, it presents comparable landmarking accuracy, despite relying on a single, random reference specimen and much sparser sampling of the structure’s surface. Second, it is performant such that it can be efficiently run on consumer-grade personal computers. Finally, it is general and can be applied to any biological structure of interest, regardless of whether anatomical atlases are available.

Our validation procedures indicate that the method is capable of recovering multivariate patterns of morphological variation that are largely indistinguishable from those obtained by manual digitization, indicating that the use of an automated landmarking approach should not result in different conclusions regarding the nature of multivariate patterns of morphological variation.

The proposed point cloud-based approach has the potential to increase the scale and reproducibility of morphometrics research. To allow ALPACA to be used out-of-the-box by users with no prior programming experience, we implemented it as a module as part of the SlicerMorph project. SlicerMorph is an extension that enables geometric morphometrics data collection and 3D specimen analysis within the open-source 3D Slicer biomedical visualization ecosystem. We expect that convenient access to this platform will make ALPACA broadly applicable within ecology and evolution.

## Introduction

In the past ten years, volumetric (3D) imaging has been used with increasing frequency to characterize morphological variation in complex biological structures in ecological and evolutionary contexts (Boyer et al., 2011; Falkingham, 2012; Marcy, Fruciano, Phillips, Mardon, and Weisbecker, 2018; Goswami et al., 2019). The general approach has been to capture high-resolution specimen images (but see (Marcy et al., 2018) and posteriorly collect the position of several anatomical landmarks of interest. These anatomical landmarks are then used in multivariate shape analyses, allowing researchers to test specific functional/developmental hypotheses regarding the ecology or evolution of complex phenotypes (e.g., Sanger et al., 2013; Sherratt, Gower, Klingenberg, and Wilkinson, 2014; Felice et al., 2019).

While the quality of imaging techniques (e.g., Gignac et al., 2016) and the density of landmarks (e.g., Goswami et al., 2019) has continuously increased during the last decade, the gold standard method for landmark data collection has remained largely the same, i.e., manual annotation by a trained expert. Manual annotation of landmarks is, however, both time and labor-intensive, low-throughput, and subject to significant amounts of intra- and inter-observer bias, precluding datasets from different laboratories (or even multi-year datasets) from being confidently combined (Percival et al., 2019).

Recently, several approaches have been developed to automate and standardize landmark data collection in the context of volumetric imaging. While deep learning approaches are starting to emerge (e.g., Devine et al., 2020), most studies approach the problem using image-based registration techniques developed in biomedical contexts (Bromiley, Schunke, Ragheb, Thacker, and Tautz, 2014; Young and Maga, 2015; Maga, Tustison, and Avants, 2017). Image registration represents the alignment of images that belong to the same anatomical structure of interest and provides researchers with a powerful framework for workflow automation, allowing morphometric research to truly enter the age of big data (Maga et al., 2017).

However, attempts to translate these biomedically-oriented approaches to more ecological and evolutionary contexts have remained rather timid and have faced substantial practical and technical barriers. For example, most image-based registration approaches depend on high-end hardware, all the while producing results in a timeframe that greatly exceeds the amount of time required for manual annotation (e.g., 10 CPU hours per specimen; Devine et al., 2020). While computing clusters have made high-end hardware more accessible at the institutional level, the cost-benefit ratio of implementing such approaches is still highly skewed against automation. Additionally, these algorithms are highly system-specific and difficult to generalize to different study systems. Finally, image registration techniques rely on specialized labor, which include a dedicated programmer for algorithmic development and an imaging technician capable of developing and troubleshooting anatomical atlases.

Here we propose a new and general approach to automated 3D landmarking based on point cloud registration. Starting with 3D surface meshes, the procedure performs pairwise registration of subjects to the specified template using a sequential procedure with four steps. Initially, the edge information in individual meshes is discarded and the resulting point clouds are downsampled in order to facilitate the initial alignment and increase the speed of calculation. These point clouds are then subjected to a global registration step (Rusu, Blodow, and Beetz, 2009), in which there is an initial alignment of the source and target point clouds. This initial alignment is followed by a local registration step (Rusinkiewicz and Levoy, 2001), in which the initial alignment is refined. Finally, the two rigidly aligned point clouds are subjected to a deformable registration step (Myronenko and Song, 2010), in which the source point cloud is deformed to match the target point cloud. As a result of the deformable registration step, the landmark correspondences across meshes gets established and landmark positions can be transferred across specimens.

Point cloud registration provides a simpler and more general alternative to image-based registration, since it not only requires less preprocessing, but is also of much lighter implementation, therefore eliminating many of the challenges listed above. Point cloud registration has three main requirements: (1) a single reference (source) specimen; (2) one or multiple target specimens; and (3) that the meshes being aligned represent the same biological structure (i.e., there are no extraneous morphological elements). Since the source specimen will be deformed to match all target specimens, some care in the choice of source mesh is advisable (e.g., avoid using individuals with extreme morphologies). However, that is not a strict requirement of the ALPACA pipeline, which allows for any individual to be chosen as the source. We provide an efficient implementation of the ALPACA pipeline in the most recent version of SlicerMorph (Rolfe et al., 2020), the 3D morphometrics extension to the open-source biomedical visualization software 3D Slicer (Fedorov et al., 2012; Kikinis, Pieper, and Vosburgh, 2014).

## Materials and Methods

Below, we present and describe in detail: (1) the set of images and landmarks used to explore the performance of the method, (2) the proposed pipeline, and (3) the metrics employed to validate the approach and to quantify how reliable it is in comparison with manual digitization and other image-based registration methods that have been published in the literature.

### Samples

When developing automated landmarking methods, it is often useful to have a dataset of manually digitized samples to serve as a reference set (i.e., a gold standard for performance). We have developed and tested our approach on a standard dataset used in many image-based automated landmarking approaches, namely the laboratory mouse skull (Maga et al., 2017; Percival et al., 2019; Devine et al., 2020). For the sake of generality, we also test it on three other datasets belonging to non-human primates (*Pongo, Pan* and *Gorilla*). Fig S1 presents the anatomical landmarks used in this study. We note, however, that our approach should work for any other biological structure of interest and it is not restricted to craniofacial research.

#### Mice

More specifically, we developed and initially tested the ALPACA framework on a published dataset containing 51 wild- and laboratory-derived inbred strains of mice (Maga et al. 2017). In short, one 8-week-old female each derived from a total of 25 inbred and 5 F1 crosses were commercially acquired from Jackson Laboratories and then sacrificed at 56±3 days of age via CO2 asphyxiation followed by decapitation. Heads were imaged using a Skyscan 1076C micro-CT using a standardized imaging protocol. These images were then processed following Maga et al. (2017). All animal procedures used in the study were reviewed and approved by the Institutional Animal Care and Use Committee of the Seattle Children’s Research Institute (protocol 13733).

#### Hominoids

We also applied the ALPACA framework to skull meshes belonging to three other mammalian species, all of them great apes: *Pan troglodytes, Gorilla gorilla*, and *Pongo pygmaeus*. These meshes were generated from CT scans of dry crania of specimens housed in the National Museum of Natural History (NMNH). We present the list of specimens used in this study as Table S1. More details can be found on Rolfe and Maga (2020).

### Pipeline - ALPACA

We approach the problem of automated 3D landmarking using a lightweight point cloud registration approach based on surface meshes. In this approach, a reference mesh (here, the source mesh) is aligned and posteriorly deformed to match a target mesh for which we want to predict the landmark positions for. Using the transformation parameters used to deform one mesh into another, we project the landmark positions of the source mesh into the target one. In other words, we approach the problem of automated landmarking by transferring the landmark position of a single specimen (or template) into another (Fig 1).

**Fig. 1.**
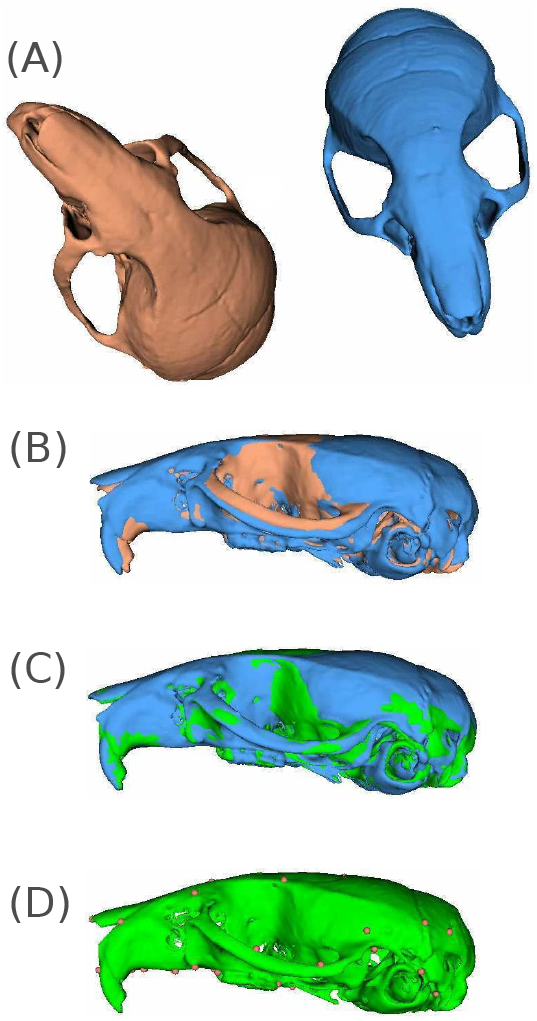
Visual representation of the ALPACA pipeline. Starting with a source (red) and target (blue) meshes representing the same biological structure and lying at arbitrary positions in XYZ axes (A), the pipeline starts with an initial downsampling of the two meshes into point clouds that are then rigidly aligned (B) to each other. Note the differences between the two meshes in terms of the angle of the nasal bone relative to the neurocranium and in terms of the position of the zygomatic arch. Once rigidly aligned, these point clouds are then subjected to a deformable registration step (C) in which the source mesh is warped (green) to match the target one (blue). Note how the nasal bone and zygomatic arch are much more closely aligned. Finally, after the deformable registration step, the landmark positions (dots) in the source mesh are projected into the target one (D) using point correspondence.

Note that the source sample does not necessarily need to be roughly aligned to the target one. Similarly, one could choose the surface mesh of any specimen for which landmark data is available as the source mesh. The same is true for the target sample. However, it is advisable to carefully consider which specimen should be chosen as the reference specimen to annotate new samples with, as the template choice may itself influence the quality of the prediction (Young and Maga, 2015). This is particularly true when the reference sample is not near the species/population average shape. The speed and ease of ALPACA pipeline is meant to greatly facilitate an initial exploration of the automation parameters, including the choice of template specimen. The user has the ability to quickly change the source sample and compare how resultant landmarks differ across samples. Based on the two initial samples, the pipeline then proceeds as follows below.

#### Step 1 - Scaling and Downsampling

The ALPACA pipeline starts with an optional step. In this step, the source mesh is isotropically scaled to match the target mesh. This step is only necessary in cases where there are large differences in size between the source and target meshes. Following the optional scaling procedure, the source and target meshes are then uniformly downsampled to simplify the initial alignment and increase the speed of calculation. The downsampling procedure occurs in units of physical space and discards the edge information, transforming the mesh into a point cloud. In our case, we downsampled each mesh using a voxel grid (Zhou, Park, and Koltun, 2018) to a point cloud of approximately 5,000 points. This value was empirically determined to be a good compromise between accuracy and computational burden (Fig 2) based on an initial exploration of hyperparameters using the mouse dataset. Once downsampled, the two point clouds are then subjected to a global registration procedure that aligns these two samples in physical space (see Fig 1).

**Fig. 2.**
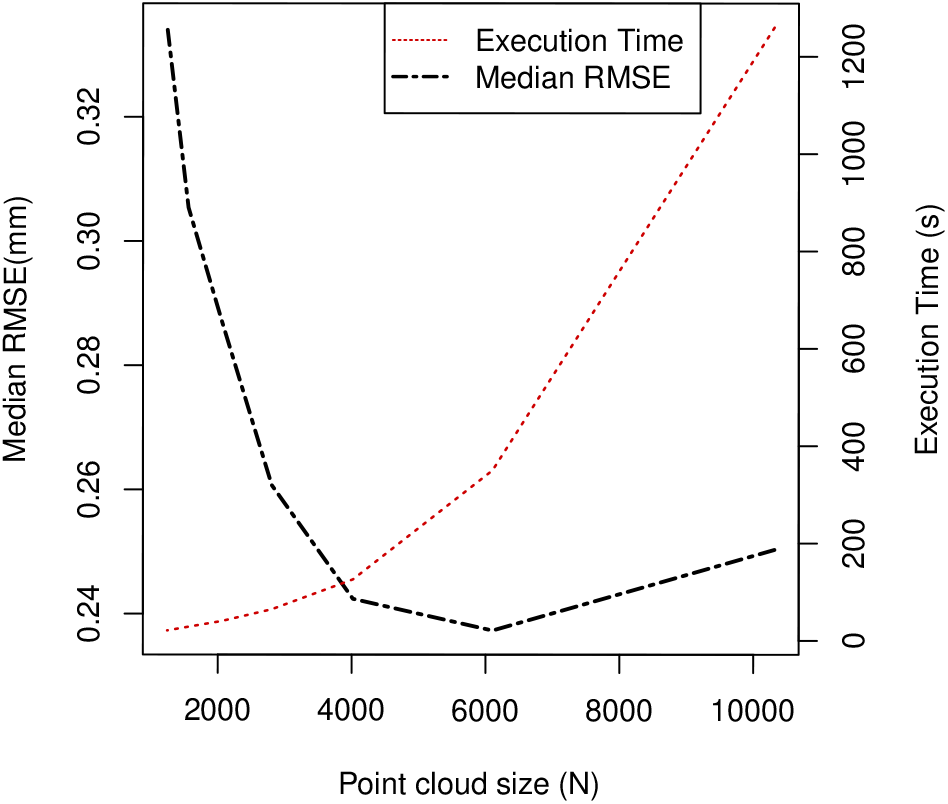
Prediction error (black dashed line) and execution time (red dotted line) of the ALPACA pipeline at different point cloud sizes. Note that prediction error, measured in terms of the root mean squared error, has a non-linear relationship with point cloud size and is minimized on point clouds in the 5,000-6,000 points range. Execution time, measured in seconds, is also non-linearly related to point cloud size, increasing exponentially with increasing point cloud size.

#### Step 2 - Global registration

Global registration techniques aim to find the optimal geometric alignment of two 3D shapes starting from initial positions (Gelfand, Mitra, Guibas, and Pottmann, 2005). In other words, it aims to find a transformation that correctly registers a source point cloud or image to a target one. Usually, approaches to global registration use iterative procedures that either (1) exhaustively search hyperparameter space for optimal combinations of parameters, or (2) that run until some convergence criteria is met (Zhou, Park, and Koltun, 2016). As such, they are often plagued by the curse of dimensionality, making them inefficient and more error prone at increasing image sizes. These approaches, however, perform well when using semantically-rich local geometric descriptors, which greatly decrease the false correspondence rate when comparing two point clouds (Chen et al., 2019). In our case, we used a feature-based random sample consensus algorithm (RANSAC) to estimate the optimal transform (Rusu et al., 2009). Essentially, in each RANSAC iteration, a user-defined number of points are sampled from the source point cloud and their corresponding points in the target point cloud are identified through finding their nearest neighbor in a space of geometric features. The features, in this case, are fast point feature histograms (FPFH) (Rusu et al., 2009). FPFH are 33-dimensional vectors that provide a description of the local geometric properties around a point and that are scale invariant, providing the RANSAC algorithm with good discriminative power in the search for point correspondence across point clouds.

Aside from the number of iterations and the number of validation steps typical of iterative algorithms, ALPACA’s implementation of the FPFH-based RANSAC has three main parameters that can be defined by the user. The first one is the normal search radius that defines the neighborhood of points used when calculating the surface normals in each point cloud. The second is FPFH search radius, which defines the neighborhood of points used when computing the FPFH features. Lastly, users have the option of controlling the maximum distance between two points up to which the points can be considered corresponding to each other.

Once the rigid transforms are obtained from the FPFH-based RANSAC, they are then fed to the third step of our pipeline, representing a local registration step.

#### Step 3 - Local registration

While the FPFH-based RANSAC algorithm provides an approximation of the rigid transform, its alignment is performed using broad geometric features, which lead to a rough alignment. In order to refine the initial alignment, we then proceeded with a local registration algorithm.

Given the initial transform obtained in the global registration step, the alignment of the two point clouds was improved upon through an iterative procedure that assigns to each point in the source point cloud to its closest point in the target point cloud. In our case, we opted using the point-to-plane iterative closest point algorithm (point-to-plane ICP) in order to do so (Rusinkiewicz and Levoy, 2001). ICP represents a family of widely used local registration algorithms, with applications in a variety of computer vision problems (Rusinkiewicz and Levoy, 2001). Essentially, given the target and source point clouds, the ICP algorithm calculates, at each iteration, the squared distance between each source point and the tangent plane at its corresponding target point. The algorithm proceeds iteratively until the distance between the two point clouds is minimized. The main advantage of the point-to-plane version of ICP is the speed of convergence relative to point-to-point error (Rusinkiewicz and Levoy, 2001).

Similar to the global registration step, the result of the point-to-plane ICP is a rigid transform, corresponding to the optional rigid alignment of the source and target point clouds. Also similar to the global registration step, users have the option to control a maximum distance between points up until which they still can be considered corresponding to each other.

Once the optimal rigid alignment is obtained, we then proceed to the deformable registration step of the pipeline.

#### Step 4 - Deformable registration and point projection

The final step of the ALPACA pipeline is the deformation of the rigidly aligned source point cloud to match the target point cloud. We use the Coherent Point Drift (CPD) algorithm in order to do so (Myronenko and Song, 2010). CPD is a probabilistic procedure for point cloud registration, in which the alignment of two point clouds is framed in terms of a probability density estimation. The source point cloud represents Gaussian mixed model (GMM) centroids that are fitted to the target set using maximum likelihood (Myronenko and Song, 2010). The main benefit of CPD is that it imposes a constraint in the form of motion coherence among neighboring points, leading to deformations that preserve the topology of the structure of interest. It also makes no other underlying assumption about the nature of the transformation itself, allowing for a wealth of possible deformation models to be true. Finally, it has the additional benefit of being one of the few methods for non-rigid registration that can accommodate large point clouds (5,000+ points) with slightly different numbers of points (Myronenko and Song, 2010).

ALPACA’s implementation of CPD contains two free parameters. Parameter β refers to the width of the Gaussian filter used when applying smoothness constraints, representing one approach to regularization. High values of beta create large directional correlation among the displacement vectors of neighboring points during deformation, and vice versa (Hirose, 2020). Parameter α, on the other hand, represents a trade-off between goodness-of-fit and model regularization, with high values leading to overall structural rigidity and low values to more structural fluidity (Hirose, 2020).

Following the deformation, ALPACA has a final and optional post-processing step in which the predicted landmarks are projected to the target surface mesh. This step guarantees that the landmarks will lay on the most exterior surface of the original mesh, limited by a user-adjustable point displacement (the default being 1% of the diagonal size of the image). Each point is projected from the deformed model to the original surface using the following steps: 1. cast a ray from the semi-landmark point on the deformed model in the direction of the normal vector at that point. Select the final intersection with the original model as the intersection point. 2. If there is no intersection, reverse the direction of the normal vector and select the first intersection with the original model. 3. In the case no intersection is found, select the closest point on the original model.

### Prediction parameters

The parameters used when running ALPACA on all four datasets are present in Table 1. When running the pipeline for the mouse dataset, we have used the synthetic population template presented in Maga et al. (2017) as the initial source mesh. While the ALPACA pipeline does not require a synthetic source template, by using the same template used for manual landmarking, we are able to directly compare the results obtained by both approaches.

**Table 1.**
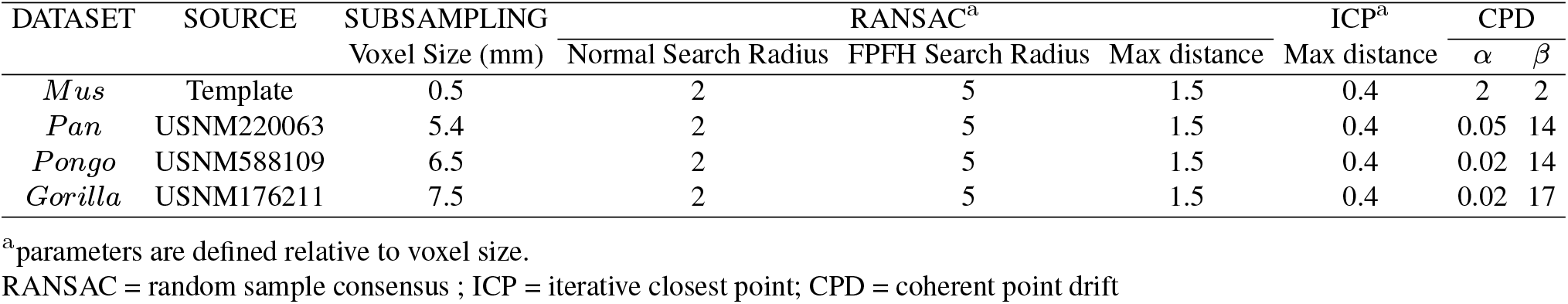
Point-cloud registration parameters used for each dataset. Parameters are divided according to the corresponding step of the pipeline. See main text for details.

When running the pipeline on the three non-human primate datasets, a randomly chosen specimen was used to generate the predictions for the remaining ones. Specimens with missing skull elements (e.g., teeth) or damaged skulls were not added to the pool of specimens from which random samples were drawn, as the algorithm assumes the presence of corresponding structures in the source and target meshes. For all four datasets, we skipped the scaling and point projection steps, as those were not necessary for these four datasets.

### Evaluating performance

We evaluated the performance of ALPACA’s approach not only in terms of the Euclidean distance between the manual and predicted landmark locations, but also in terms of the patterns of morphological variation and covariation among landmarks.

#### Euclidean distance

One way in which automated landmarking datasets can be evaluated is through the calculation of the root mean squared error (RMSE) between the landmark positions as predicted by the pipeline and as manually annotated (e.g., Percival et al., 2019). RMSE values calculated for such datasets include not only the errors committed by the automated pipeline but also observation errors committed by the anatomical expert. While the pipeline error can be minimized, intra-observer error is unavoidable. As a consequence, the error produced by the pipeline should always be evaluated on a relative basis (Percival et al., 2019). We here consider the intra-observer error as the minimum possible error the approach could hope to achieve and, therefore, use it to evaluate relative performance. While measurements of intra-observer error are difficult to be obtained for most datasets, the mouse skull and mandible have become a standard dataset in many automated landmarking algorithms (e.g., Young and Maga, 2015; Maga et al., 2017; Percival et al., 2019; Devine et al., 2020). As a consequence, there are precise published estimates of intra-observer error for the majority of our landmarks (35 of 45; Percival et al., 2019). We here assume that the intra-observer manual annotation errors reported by Percival et al. (2019) are, to a large extent, representative of the morphometric community at large.

Since the non-human primate datasets have not been measured repeatedly, we report their overall RMSE values but evaluate these datasets purely in terms of their ability to accurately characterize size and shape variation.

#### Size and Shape

In order to quantify the impact of the choice of method on the results of size and shape analyses, we performed a joint Procrustes superimposition across datasets (Rohlf and Slice, 1990). We then performed a Procrustes MANOVA with landmarking method as a factor (Adams, Collyer, Kaliontzopoulou, and Sherratt, 2016), leading us to quantify the percentage of the total variance in shape that is associated with the choice of landmarking methodology.

We also use the joint superimposition to test whether Procrustes distances obtained by each method are significantly different from one another using a paired signed Wilcoxon rank test (Wilcoxon, 1992).

To evaluate the ordination of specimens in size and shape space, we performed principal component analyses of the tangent space coordinates and correlated the manual predictions with the automated ones in terms of the centroid size and principal component scores following Maga et al. (2017).

Finally, to evaluate the similarity in the relative distribution of morphological variation in multivariate space, we compare the percent variance explained by each principal component across both methods and test whether observed differences are larger than differences that should be expected solely based on sampling error. The rationale used when comparing measurement error to sampling error is simple. Automated methods allow for substantial increases in the sample sizes of most studies (e.g., Porto and Voje, 2020). As a consequence, the smaller the error produced by the pipeline relative to sampling error, the less significant measurement error becomes in terms of its effect on the study’s outcome.

### Bias correction

In many situations, researchers might be interested in combining datasets generated by automated landmarking methods with manually annotated ones (Percival et al., 2019). That is often a challenging task, since there is the possibility that both the means and error variances are different across landmarking methods. Hence, when possible, it is generally not advisable to do so. However, there are a few situations in which such interest might be justified. For example, a researcher might be interested in combining multi-year (large) datasets that were acquired using different methods by a laboratory producing advanced intercross lines (e.g., Cheverud et al., 2014). In that case, we here propose the usage of a parametric empirical Bayes framework (Fortin et al., 2018) to robustly adjust the tangent space coordinates for these effects. The ComBat model assumes that the expected values of the tangent space coordinates can be modeled as linearly dependent on (landmarking) method-specific effects, and whose errors are also method-specific. The underlying assumption is that the landmarking method has both additive and multiplicative effects on the tangent space coordinates (Fortin et al., 2018). The outputs of this linear model represent the bias-corrected tangent space coordinates, if we assume the manual landmarking method to be the reference one.

In our case, the only dataset large enough to apply bias correction was the mouse dataset. In order to do so, we selected a small percentage of the original samples (20%) to estimate the ComBat model parameters and use such parameters to correct the automated predictions for all remaining samples (80%), We then evaluate the impact that such procedure has on RMSE estimates and overall mean configuration plots.

### Implementation - SlicerMorph module

All algorithms were implemented as a SlicerMorph module using the following external python libraries: *open3d v.0.9.0* (Zhou et al., 2018) and *pycpd 2.0.0*. Source code for SlicerMorph can be found at https://github.com/SlicerMorph/SlicerMorph or downloaded directly from the 3D Slicer version 4.11.0 Extension Manager. The ALPACA module provides a graphical user interface and can be run on any operating system that Slicer runs. As Supporting Information, we provide installation instructions and links to a detailed ALPACA tutorial. The ALPACA pipeline was implemented with two different modes of functionality: a pairwise and a batch-processing. The pairwise branch should largely be used to perform a user-guided search for the best combination of parameters for their dataset, which can then be applied to a larger array of samples in batch mode. Note, however, that SlicerMorph module was developed with a larger array of use cases than the four datasets we present here and, therefore, presents default parameters that might not be ideal for all study systems. Similarly, largely due to the underlying RANSAC implementation, ALPACA is not deterministic by necessity. In the mouse dataset, repeated runs using the same specimens and the same hyperparameters will produce final predictions that are 0.03 mm apart, on average.

## Results

### Implementation speed

We used the mouse dataset (N=51) to benchmark the speed of the pipeline, using a Linux Mint OS laptop with an Intel Core i7-6700HQ 2.7Ghz CPU and with 16GB of RAM. When run multi-threaded, the complete ALPACA workflow for 51 samples took approximately 1.5 h. The majority of the time (>50%) was spent on the deformable registration step. When run pairwise, the breakdown per specimen (on average) is as follows: 0.67s for downsampling, 1.42s for global registration, 8.27s for local registration (2.23s of which are devoted to the calculation of surface normals), and 287.18s for deformable registration.

### Manual landmarks vs. ALPACA landmarks

#### Euclidean distances

In terms of their Euclidean distances to manual landmarks, which we treat as gold-standard, the majority of ALPACA’s landmarks were accurately placed by the pipeline. In the mouse dataset, median RMSE values varied from 0.16 mm to 0.64 mm (Fig 3A), with an average of 0.23 mm. These RMSE values are largely comparable to those obtained by other image-based registration methods (Fig 3, blue dots, Percival et al., 2019) and also close to the manual (intra-observer) error (Fig 3, green squares). Note that intra-observer error represents the theoretical minimum error the pipeline could hope to achieve. Only landmarks 2 and 3 present much higher than average error when compared to the manual dataset and those are associated with a methodological bias in the estimate of the population mean, as it will be further explored in the Procrustes analysis section.

**Fig. 3.**
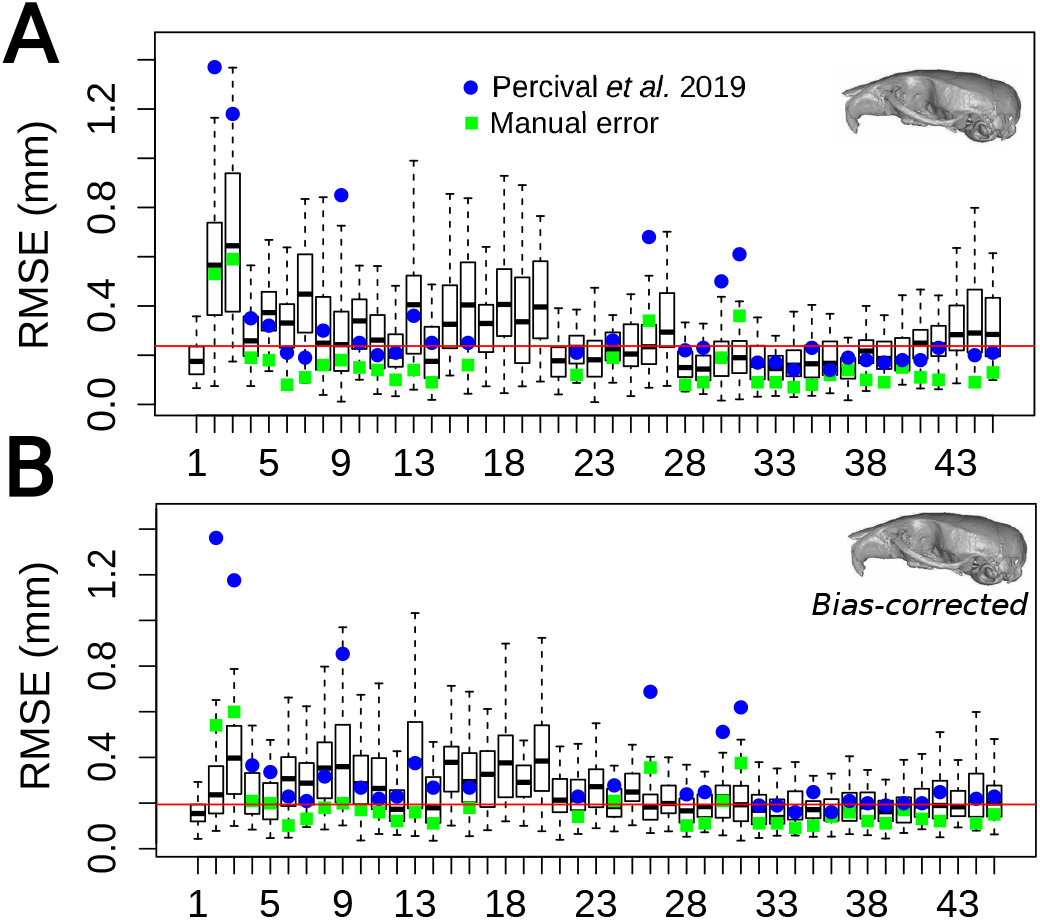
Boxplots illustrating the landmark-specific distribution of prediction errors for the mouse dataset, measured as root mean squared error (RMSE). (A) Initial predictions. (B) Bias-corrected predictions. RMSE values are calculated based on the difference between the predicted landmark positions and their manually annotated counterparts (in mm). The median error across landmarks is illustrated with a red line. Prediction errors calculated for an image-based registration approach are illustrated with blue circles (Percival et al., 2019). Intra-observer errors calculated by Percival et al. (2019) are presented as green squares.

Bias-correction considerably reduced mouse RMSE values for the majority of landmarks and effectively corrects it for the biases on the population means (Fig 3B). On average, mouse RMSE values after correction are 0.04 mm lower than the uncorrected ones (Fig 3B), resulting in a mean value of 0.19 mm. Finally, we should also highlight that the maximum improvement that could be expected in any future automated algorithm is in the order of 0.03 mm, given the difference between bias-corrected RMSE values (0.19 mm) and manual RMSE values (0.16 mm).

Among the great apes, median RMSE values were generally more homogeneous across landmarks and largely comparable across species relative to their overall skull size (Fig 4). *Gorilla* median RMSE values varied from 3.55 mm to 5.36 mm, with an average of 3.89 mm. *Pan* median RMSE values varied from 2.86 mm to 4.35 mm, with an average of 3.06 mm. Finally, *Pongo* median RMSE values varied from 3.92 mm to 7.18 mm, with an average of 4.35 mm.

**Fig. 4.**
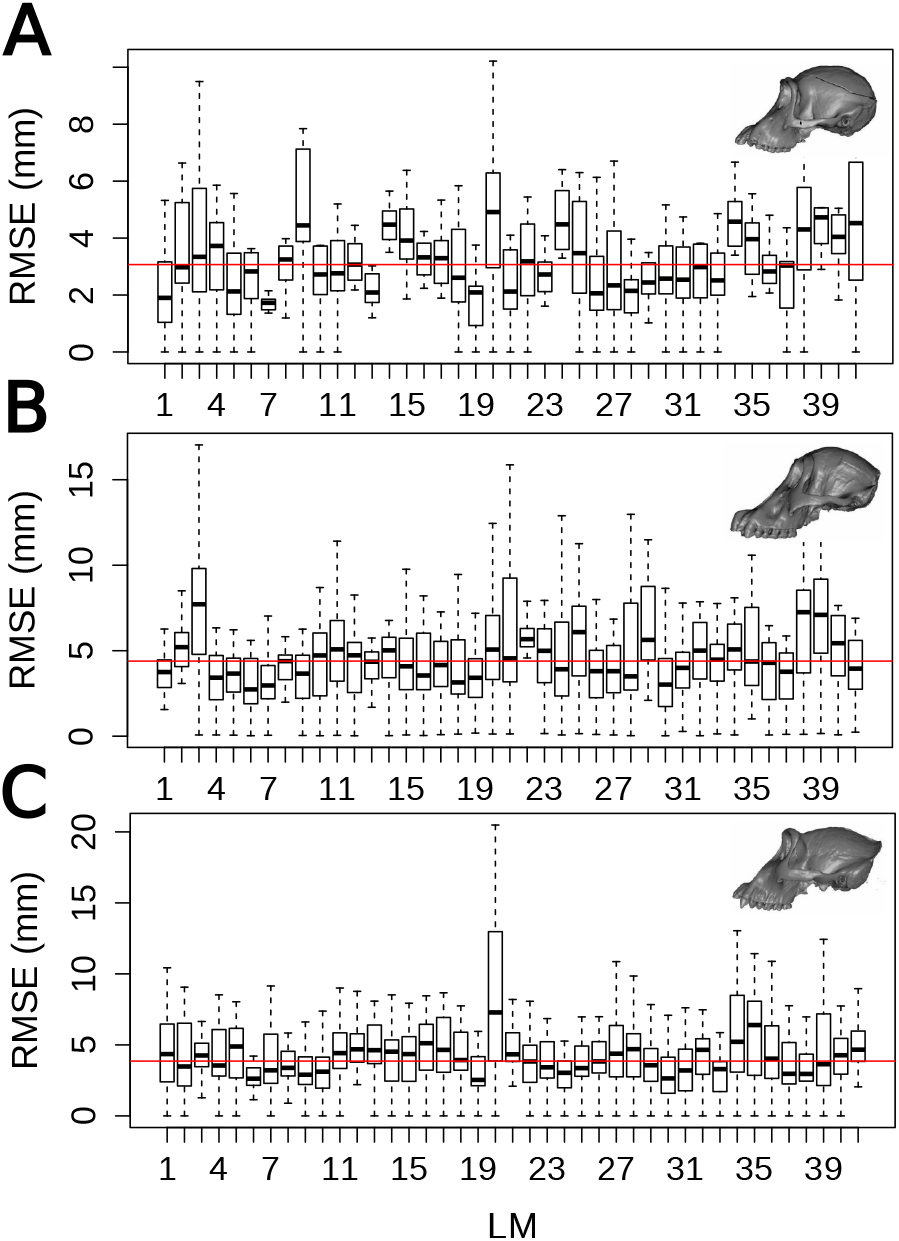
Boxplots illustrating the landmark-specific distribution of prediction errors for all three great ape datasets, measured as root mean squared error (RMSE). RMSE values are calculated here based on the difference between the predicted landmark positions and their manually annotated counterparts (in mm). The median error across landmarks is illustrated with a red line. (A) *Pan*; (B) *Pongo*; (C) *Gorilla*.

When standardized by the maximum skull length, mean RMSE values observed across species are equivalent to: 1% (*Mus*), 1.18% (*Gorilla*), 1.57%(Pan), 1.81% (Pongo), indicating similar performance of ALPACA across different organisms.

#### Procrustes analysis

In Fig 5, we report a joint generalized Procrustes analysis (jointGPA) of manual- and ALPACA-based landmark datasets. When compared to the manually digitized landmarks, the ALPACA-based landmarks show a remarkably similar scatter after a joint superimposition (Fig 5A, D-F). On the mouse dataset, the manually digitized dataset possesses a wider spread around the mean shape than ALPACA dataset, as revealed by a paired signed Wilcoxon rank test (p<0.001). This difference is particularly noticeable for landmarks in the lateral parts of the nasal/premaxilla sutures (Fig 5A, landmarks 2 and 3) and is effectively removed by the bias-correction procedure (Fig 5C, p=0.83). No other dataset presents significant differences in the degree of shape variation (p=0.86 for *Pan*, p= 0.88 for *Pongo*, and p=0.177 for *Gorilla*).

**Fig. 5.**
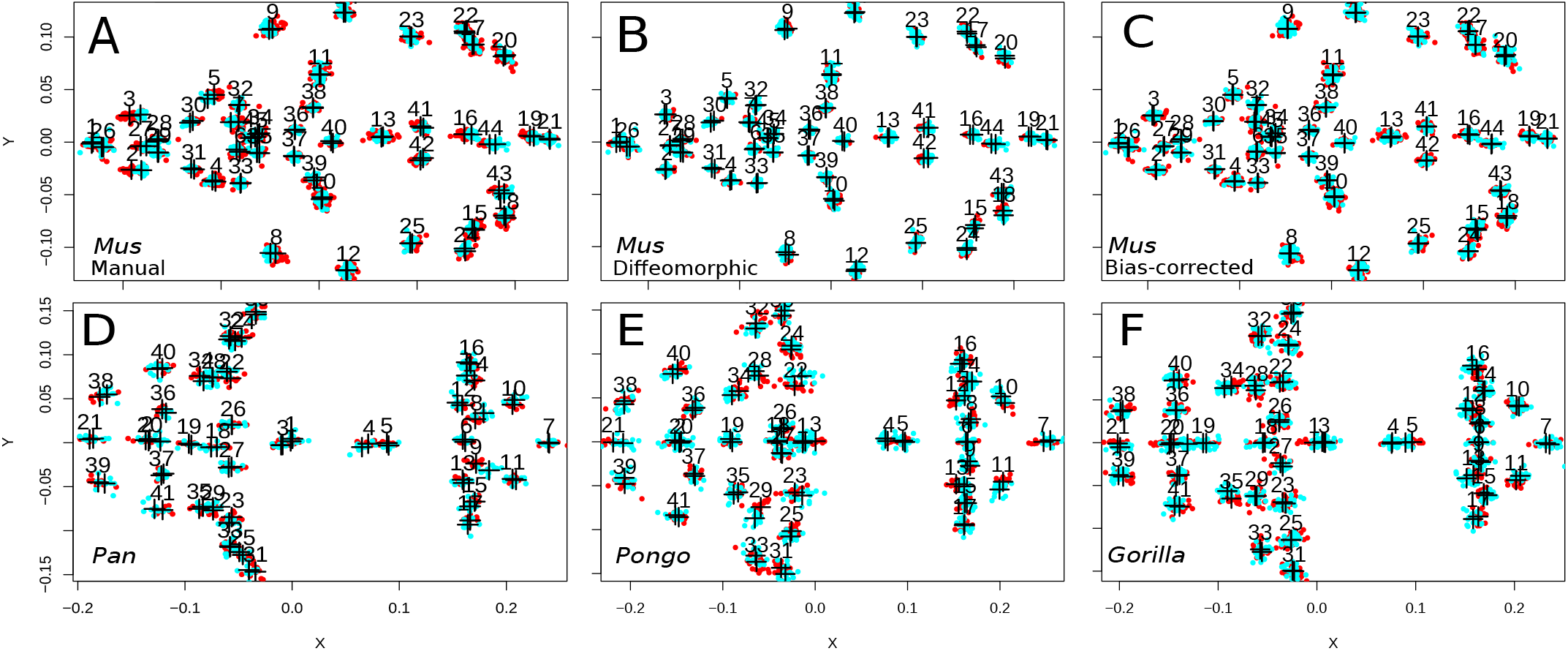
Two-dimensional projection (XY) comparing theALPACA landmark predictions (light blue) with other methods (red) afterjointGPAsuperimposition. Crosshairs indicate the consensus shapes of each method underthe joint superimposition. Other methods are, in order: (A) Manual landmarking (Mus); (B) Diffeomorphic approach described in Maga et al. (2017) (Mus); (C) Manual landmarking after bias correction (Mus); (D) Manual landmarking (Pan); (E) Manual landmarking (*Pongo);* and (F) Manual landmarking (Gorilla).

A Procrustes multivariate analysis of variance (procMANOVA) conducted in the jointGPA data reveals that the landmark placement method (ALPACA vs Manual) explains around 14.6% of the total variation around the mean shape in the mouse dataset (p<0.001). After bias correction, the landmark placement method loses its explanatory power (1%). Landmark placement method explains comparable percentages of variation in all three ape datasets (16% for Pan, 16.2 % for *Pongo*, and 8.2% for *Gorilla*).

Despite the difference in the overall magnitude of variation around the mean shape in the mouse dataset, the distribution of morphological variation in multivariate space is virtually identical between the two approaches in all four datasets (Fig 6). Observed differences fall well within the 95% bootstrap resamples of their respective dataset, indicating that the amount of error generated by the pipeline is substantially lower than the expected sampling error at their corresponding sample sizes.

**Fig. 6.**
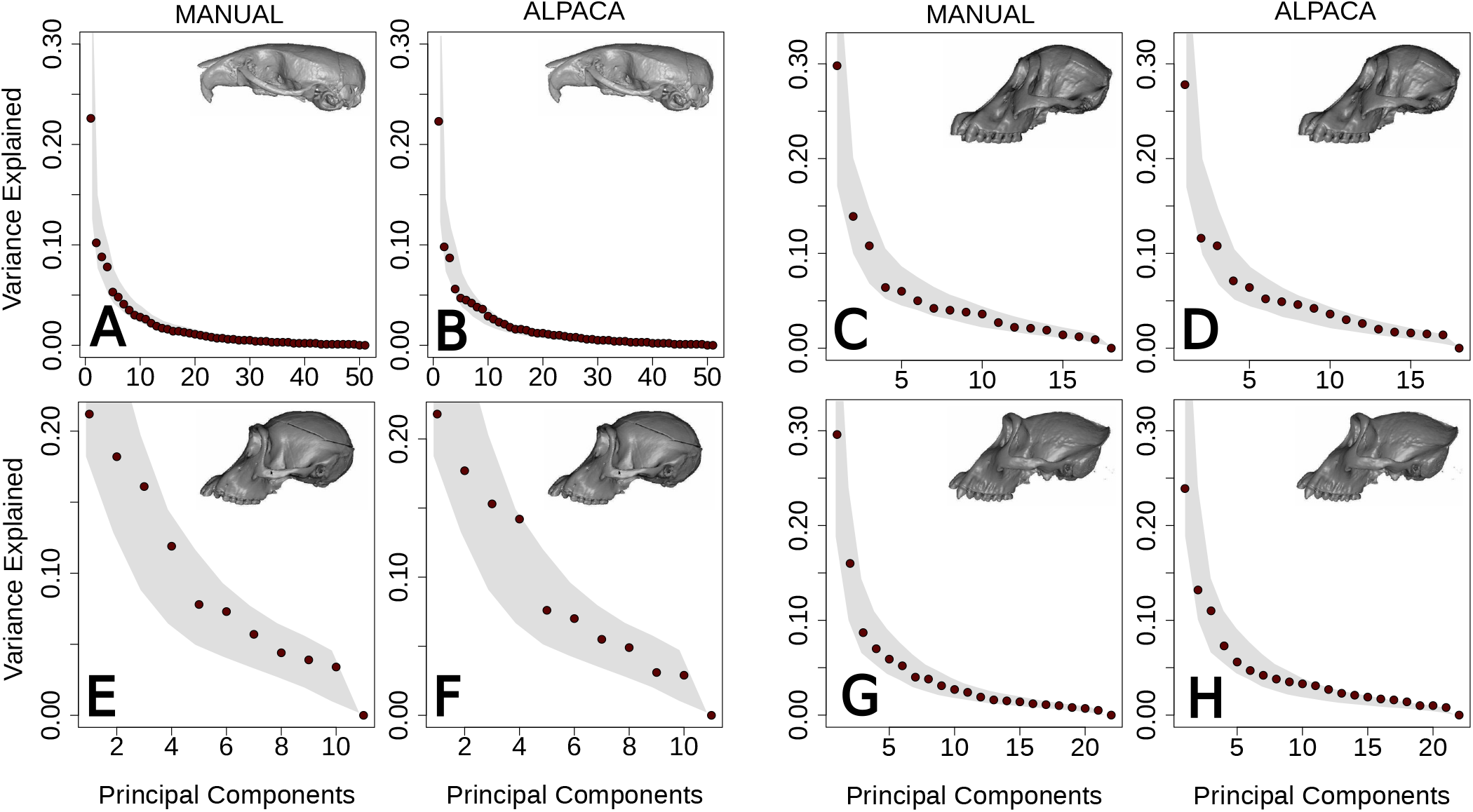
Percent variance explained by each principal component across different PCAs. ALPACA predictions are presented to the right of the manual results. Shaded areas in both plots indicate the 95% confidence interval of the eigenvector variances in the manually annotated dataset. (A-B) Mus; (C-D) *Pongo*; (E-F) Pan; (G-H) *Gorilla*.

Similarly, both the centroid sizes and shape PC1 scores for each specimen are also highly correlated across methods (Fig 7). Other PCs show similar patterns. Correlation coefficients for the remaining PCs are provided as Supporting Information Table S2.

**Fig. 7.**
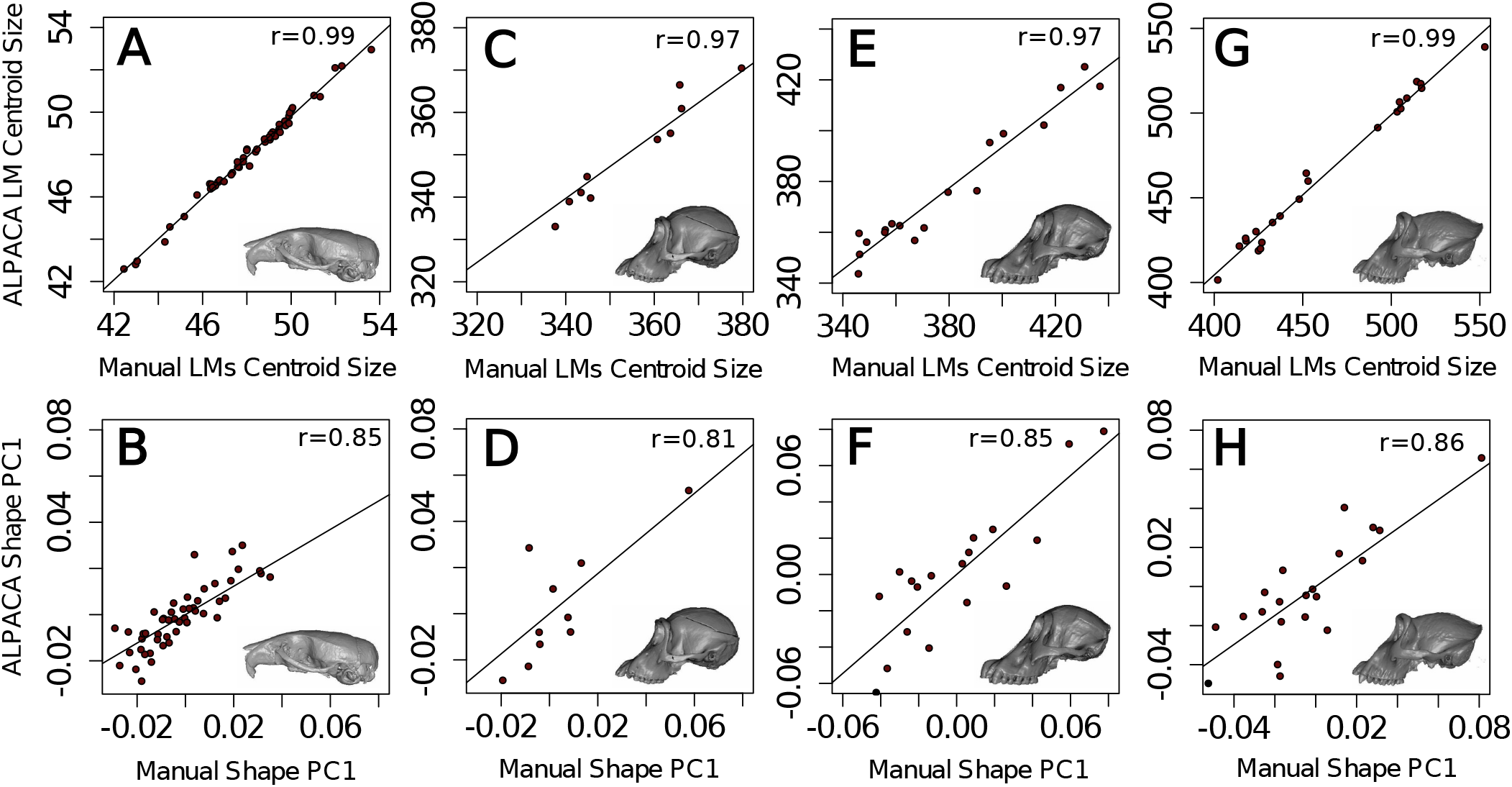
Comparison of centroid size (A-C-E-G) and shape PC1 (B-D-F-H) measures as predicted by ALPACA and by manual annotation. Note the high and statistically significant correlations (p<0.001) across all datasets. Correlation values for the remaining PCs are presented in Table S2. (A-B) *Mus*; (C-D) Pan; (E-F) *Pongo*; (G-H) *Gorilla*.

### ALPACA landmarks vs. Diffeomorphic landmarks

Since the mouse dataset reported in this study has been used to develop an image-registration pipeline (Maga et al., 2017), we also report a joint GPA comparing ALPACA to more traditional image-registration workflows.

The ALPACA-based landmarks show a virtually identical scatter when compared to the diffeomorphic ones after a joint superimposition (Fig 5B). Both methods also possess similar spreads around the mean shape. A procMANOVA conducted in the jointGPA reveals that there is a significant difference between the multivariate means obtained for both datasets (p<0.001). However, contrary to manual datasets, a paired signed Wilcoxon rank test indicates that ALPACA specimens have, on average, the same magnitude of procrustes distances compared to the diffeomorphic methods (p = 0.997). Not surprisingly, the distribution of morphological variation in multivariate space is also similar between the two approaches (Fig 6; Fig 8), with ALPACA having a more similar distribution to the manually annotated dataset (Fig 6). Finally, both the centroid sizes and shape PC1 scores for each specimen are also highly correlated across methods (Fig 8). Other PCs show similar patterns. Correlation coefficients for the remaining PCs are provided as Supporting Information Table S2.

**Fig. 8.**
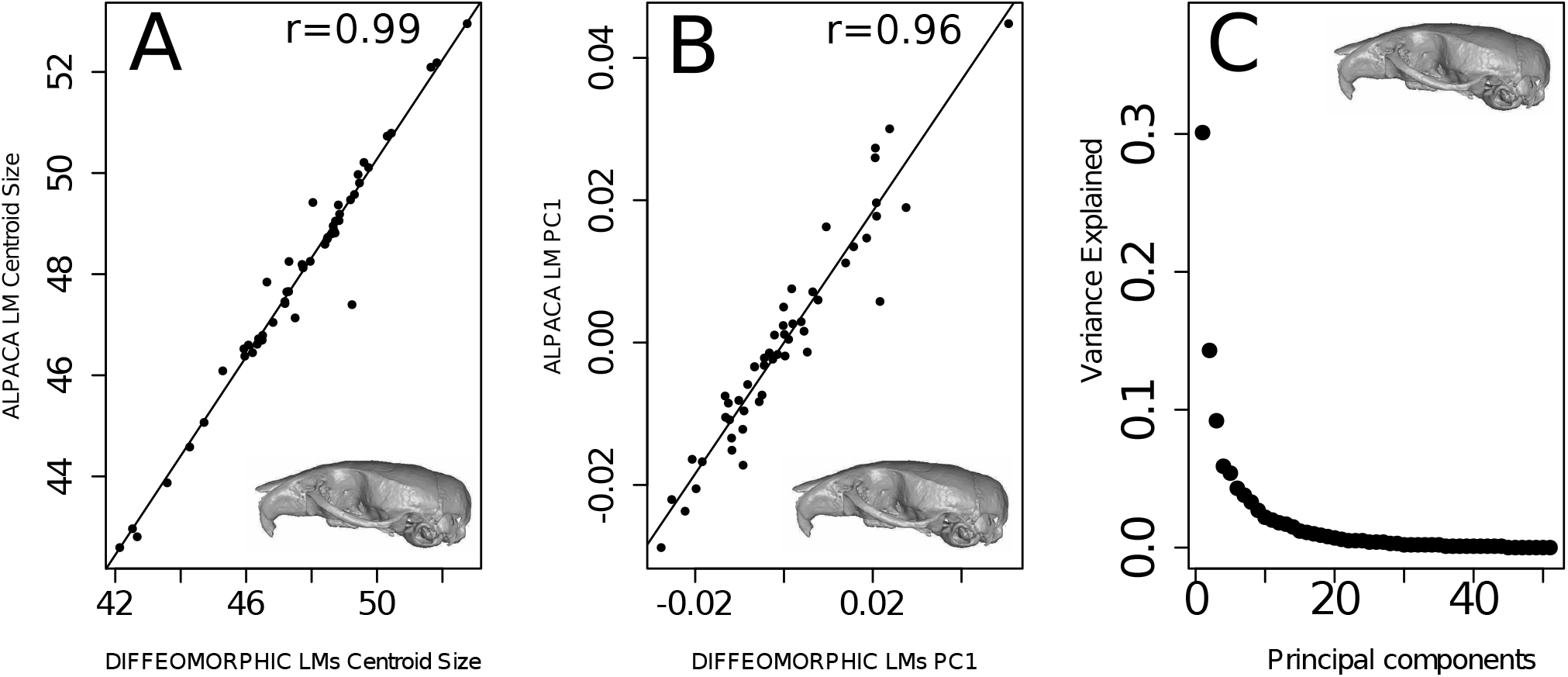
Comparison of centroid size (A) and shape PC1 (B) measures as predicted by ALPACA and by the diffeomorphic method of Maga et al. (2017) based on the mouse dataset. Note the high and statistically significant correlations (p<0.001) between the two methods. Correlation values for the remaining PCs are presented in Table S2. (C) Percent variance explained by each principal component according to the diffeomorphic method. ALPACA results are shown in Fig 6.

## Discussion

Dense morphometric characterizations of biological structures have become an essential component of morphological studies in ecology and evolution (Souter, Cornette, Pedraza, Hutchinson, and Baylac, 2010; Bardua, Felice, Watanabe, Fabre, and Goswami, 2019; Collins, Edie, Gao, Bieler, and Jablonski, 2019; Goswami et al., 2019). As a consequence, the gold standard for morphometric data collection (i.e., manual digitization) has become an important bottleneck for morphometric research pipelines. Here, we propose a fast and accurate pipeline for automated landmarking in any three-dimensional biological structure. Our approach is based on a lightweight point cloud registration approach, which can be used to transfer landmarks from a single source specimen to one or multiple targets, accurately placing landmarks of interest on unmeasured specimens.

### ALPACA advantages

One main benefit of our approach relative to other automated methods is that it works with a single reference specimen, making it more broadly applicable to ecological and evolutionary research. While anatomical atlases are available for mouse datasets (e.g., Maga et al., 2017), they are rare for non-model organisms and represent an important constraint for image-based approaches. Other advantages of our framework are its accuracy, speed, consistency and its low hardware requirements.

#### Accuracy

When applied to the mouse dataset, our approach obtains results that are virtually identical to other imagebased registration techniques (Fig 5), and recovers patterns of morphological variation that are also indistinguishable from those obtained by manual digitization (Fig 6). While we observe greater Procrustes distances in the manually annotated mouse dataset when compared to automated one, this is a common observation in many image-registration based approaches (e.g., Boyer et al., 2011; Maga et al., 2017; Percival et al., 2019; Devine et al., 2020) and is partially explained by the intra-observer error present in manually annotated ones. As a matter of fact, we can calculate how much room still exists for methodological improvement based on the mouse dataset and this value seems to be in the order of 0.03 mm, given the small difference between bias-corrected RMSE values and manual error values (Fig 3). Furthermore, since we have produced a SlicerMorph module that has a graphical user interface, any apparent error produced by the pipeline can be instantly corrected using the 3D Slicer’s fiducial tools (Fedorov et al., 2012; Kikinis et al., 2014).

#### Speed and consistency

The other large benefit of automation is its overall speed and consistency. Calculating speed in morphometric data collection is fraught with difficulty, since both manual and automated approaches require some preprocessing steps whose speed is hard to quantify. For manual landmarking, the researcher annotating the dataset will need to spend time getting used to the order of landmarks and the overall layout of the annotation software. These steps are not necessary in automated approaches. Automated approaches will require, on the other hand, a more thorough cleaning of each sample’s mesh, due to the need for structural correspondence across samples. This cleaning step will often require segmentation of the morphological elements of interest and removal of any extraneous information (e.g., removal of neck vertebrae from a skull mesh). In other words, if sample pre-processing cannot be automated, the speed of the algorithmic pipeline can be counterbalanced by bottlenecks in sample preparation. Cleaning steps are often much simpler for manual landmarking, since the user can start from an image sequence containing a myriad of extraneous morphological elements. As such, ALPACA’s speed, as reported below, should be taken with a grain of salt.

The ALPACA pipeline performs landmark prediction for the entire mouse dataset (N=51) in 1.5 hours, when run multithreaded. Note that the number of landmarks annotated in each mouse skull is on the smaller side of current morphometric approaches (Souter, Cornette, Pedraza, Hutchinson, and Baylac, 2010; Bardua, Felice, Watanabe, Fabre, and Goswami, 2019; Collins, Edie, Gao, Bieler, and Jablonski, 2019; Goswami et al., 2019) and that ALPACA’s execution time is largely independent on the number of landmarks. Given the recent popularization of high density semi-landmark approaches in ecology and evolution (Gunz and Mitteroecker, 2013), ALPACA would allow high-density morphometric characterizations of numerous specimens in a matter of hours.

More importantly, ALPACA’s main advantage over manual annotation is its consistency. As mentioned before, manual annotation is subject to significant amounts of intra- and interobserver bias. These biases are often in the same order of magnitude as intra-specific differences (Robinson and Terhune, 2017) and represent an important and understudied issue in the field. Currently, the only way to address concerns about bias in morphometric studies is through the use of multiple expert annotators. By using ALPACA, a single template can be used by a research group throughout the years, allowing for a better standardization of landmarking protocols and increasing its reproducibility. Similarly, even multiple research groups can use the same template, allowing for data to be combined across studies from different laboratories.

#### Hardware and ease-of-use

Finally, another major advantage of our framework is the ability to obtain high-throughput phenotyping with consumer-grade, off-the-shelf hardware. As currently implemented, the SlicerMorph module can be run on any machine with Windows 7 64-bit, MacOS X Lion, or recent Linux distribution, 8GB of RAM memory, 1280×1024 monitor resolution, and graphics card with at least 1GB memory. For ease-of-use, the pipeline was implemented with two branches: pairwise and batch-processing. The pairwise branch can be used to explore and fine-tune the registration parameters by going through the process of registering a single (target) sample to its reference (source), step-by-step. This step-by-step approach allows users to find the best combination of parameters for their task, which can then be applied to a larger array of samples in batch mode. In other words, the batch-processing branch opens up the possibility of a simple automated pipeline for high-throughput high-dimensional phenotyping, which will greatly increase the scale morphometric approaches in ecology and evolution.

### ALPACA limitations

The main limitation of the proposed pipeline, which is largely shared with other deformable registration approaches, is that it can lead to spurious results when the initial shapes are too dissimilar and/or registration parameters are poorly chosen (Boyer et al., 2011; Percival et al., 2019). In other words, when working in a broad phylogenetic context, with species that are highly divergent in shape and form, the ability to find corresponding landmarks can break down. In our view, the proposed pipeline is better employed within species or among closely related species. In broad phylogenetic contexts, a more careful consideration of the source meshes will likely be necessary. One possible approach would be to use multiple source meshes, each corresponding to a clade or morphotype. In that case, ALPACA would still increase the speed and reproducibility of morphometric data collection, but it would have to be applied separately for each clade or morphotype. The application of ALPACA in this context would, as a consequence, allow researchers to sample more deeply within a clade or morphotype, thus increasing the overall sample size and improving the robustness of the morphometric results. Another possibility would be to use homology-free landmark approaches (Boyer et al., 2015). One should note, however, that homology-free approaches, such as Auto3DGM (Boyer et al., 2015), generally do not allow for the addition of samples post-hoc. In other words, if a new sample needs to be incorporated into the dataset, the pipeline has to be rerun for all samples. This limitation is not present in ALPACA, since the use of a reference specimen (or template) allows for the addition of new samples post-hoc, giving the user more flexibility.

Another limitation of the ALPACA is its sensitivity to the presence of noise in the form of additional skeletal structures or damaged parts (Myronenko and Song, 2010). In mouse datasets, for example, neck vertebrae and limbs are often still attached to the base of the skull. If skull segmentation is not properly carried out and rigidity constraints in the deformable step are not correctly fine-tuned, the addition of such skeletal elements to the 3D surface can lead to spurious results. Similarly, primate skulls frequently present missing canines/incisors and males tend to present largely developed sagittal crests. Such missing, extremely dimorphic or damaged skeletal elements can potentially lead to increased prediction error. Note, however, that several of the primate skulls used in the current manuscript do present such artifacts and, therefore, the results presented here represent the actual performance of the method even when faced with significant challenges. In other words, due to the rigidity constraints in the deformable step, damaged or missing skeletal structures can be overcome (to some extent) by the pipeline when properly tuned (Fig S2). At some point, however, the differences will be too extensive for rigidity constraints to account for them. ALPACA is not unique in this sense, these constraints (i.e., completeness, and sensitivity additional objects) are common in almost all automated morphometric analysis.

## Conclusions

We have developed a lightweight point cloud registration approach (ALPACA) for automated landmarking of three-dimensional biological structures represented by surface meshes. ALPACA greatly increases the scale and reproducibility of three-dimensional geometric morphometrics, and opens up new research avenues for morphometric research.

## ACKNOWLEDGEMENTS

This project is funded by an NSF grant (An Integrated Platform for Retrieval, Visualization and Analysis of 3D Morphology From Digital Biological Collections, award number 1759883) and by a NIH/NIDCR grant (Inbred Mice Strains: Untapped Resource For Genome-Wide Quantitative Association Study For Craniofacial Shape, DE027110) to AMM. We thank participants of the Spring 2020 SlicerMorph Workshop for their valuable feedback on initial iterations of ALPACA.

## Supporting Information 1

### Installation instructions

1. Download the 3D Slicer preview from: https://download.slicer.org/?date=2020-09-16. The preview release revision 29373 has been tested in all three main operating systems (Mac, Windows, Linux) for the purposes of this manuscript, but any recent version of 3D Slicer should work.
2. While installing, if you receive warning about security or unknown publisher, ignore and proceed with the installation. If you are a Mac user and you are getting an error message with regard to security, you might have to do these steps to run Slicer after installation:

a. Find the app on Finder
b. Right click and select “Open”
c. When it says “this app can’t be opened”, hit cancel
d. Right click and select “Open” a second time.
e. Select “Open anyway”
3. Within 3D Slicer, open the “Extension Manager”, go to the “Install Extensions” tab, and search for SlicerMorph. Click “Install”. Wait until all nine extensions are shown under the “Manage Extensions” tab, and then click “Restart”. After the restart, open the module drop-down menu and you should see that SlicerMorph is now listed.
4. Click on ALPACA (SlicerMorph->SlicerMorph Labs). This will download and install an external python library called *Open3D*. It is a large library and it may take 5-10 minutes to install, during which Slicer will look like it stalled. Be patient. Mac users who are using MacOS earlier than 10.15, need to follow this instruction in their Python console:

*pip_uninstall(‘open3d’*)
*slicer.util.pip_install(‘open3d==0.8.0’)*
5. If you haven’t encountered any issues, at this point you should be set. If you want to be absolutely sure, please type these commands to your python window to double-check:

*import open3d*
*import pycpd*
6. If you end up getting error message about any of these libraries being missing, then you can try manual installation from your python console:

*pip_install(“name_of_the_library”)*
7. A detailed ALPACA tutorial can be found at:

*https://github.com/SlicerMorph/S_2020/tree/master/Lab_ALPACA*

**Table S1:**
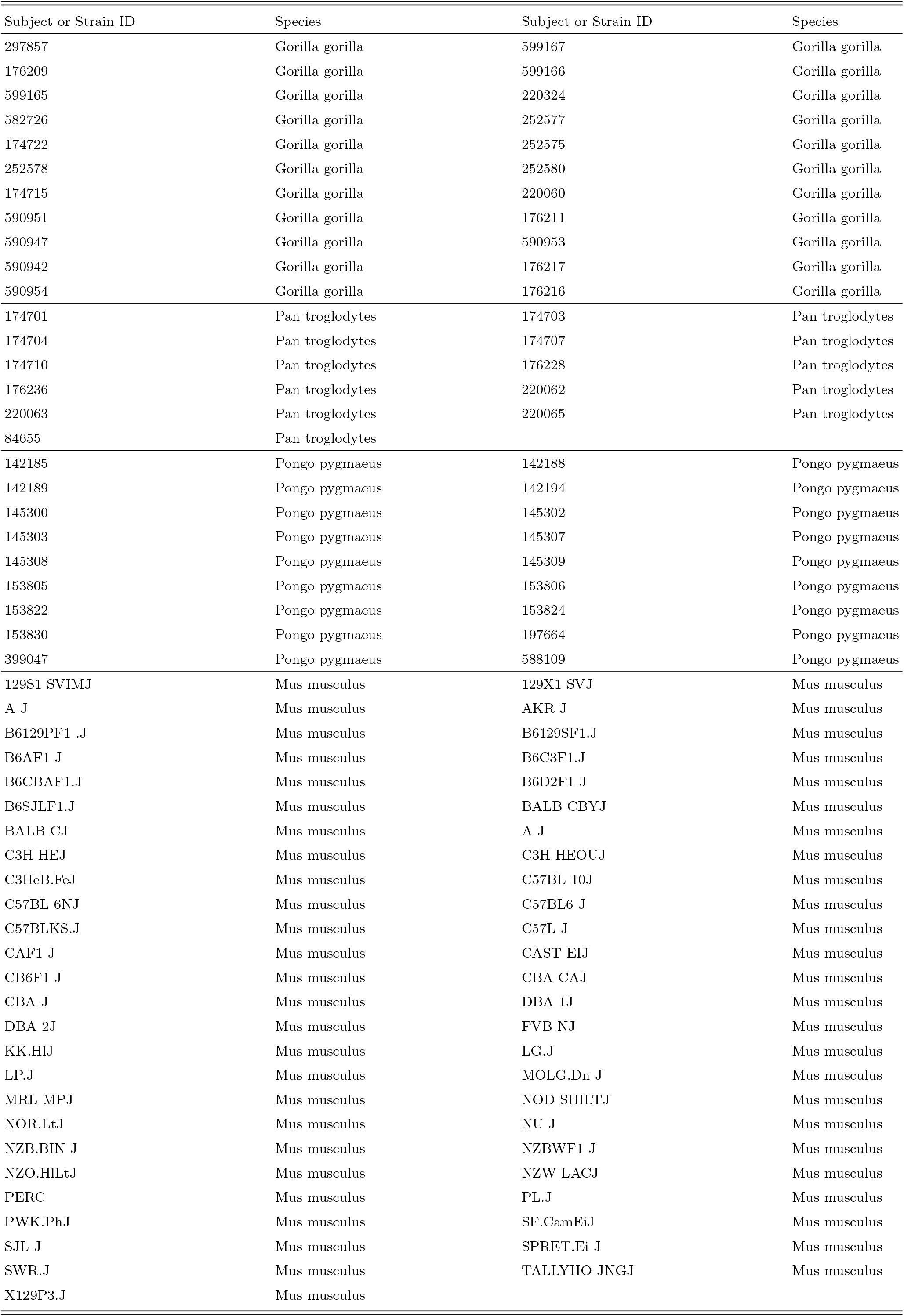
Specimens used in this study.

**Table S2:** Estimated correlation between the method-specific principal component scores (PC2-PC5) of individual specimens based on a joint Generalized Procrustes analysis.

| Variable | Manual vs ALPACA |  |  |  | ALPACA vs Diffeomorphic<sup>a</sup> |
| --- | --- | --- | --- | --- | --- |
|  | *Mus* | *Pan* | *Pongo* | *Gorilla* | *Mus* |
| PC2 | 0.78 | 0.88 | 0.78 | 0.65 | 0.85 |
| PC3 | 0.76 | 0.90 | 0.42 | 0.88 | 0.85 |
| PC4 | 0.54 | 0.92 | 0.84 | 0.54 | 0.26 |
| PC5 | 0.32 | 0.65 | 0.76 | 0.80 | 0.86 |

<sup>a</sup>Maga et al. (2017)

**Fig. S1.**
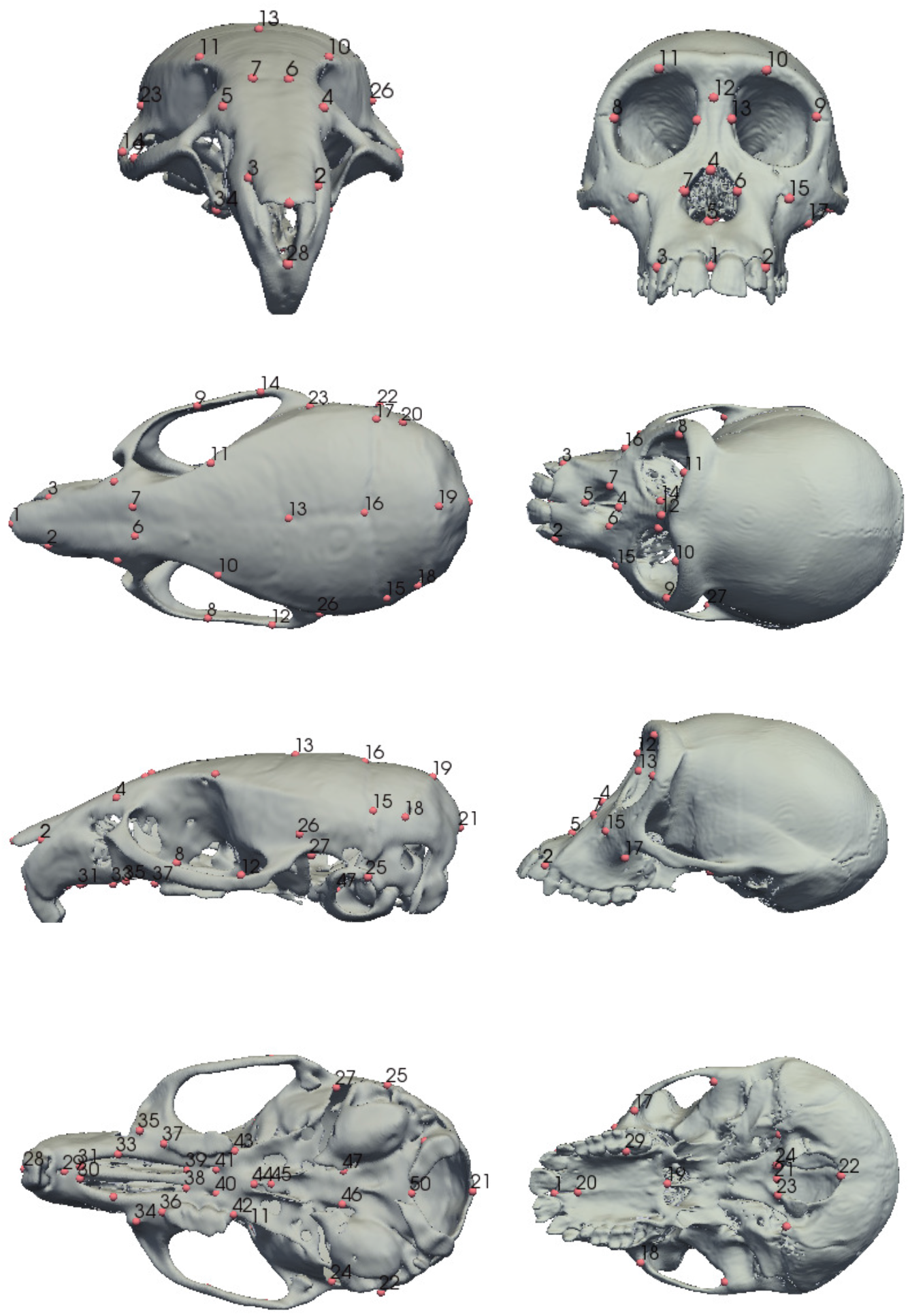
Anatomical landmarks used in morphometric analysis from frontal, superior, lateral and inferior views. First column illustrates landmarks collected in each mouse skull. Second column illustrates landmarks collected in each hominoid skull.

**Fig. S2.**
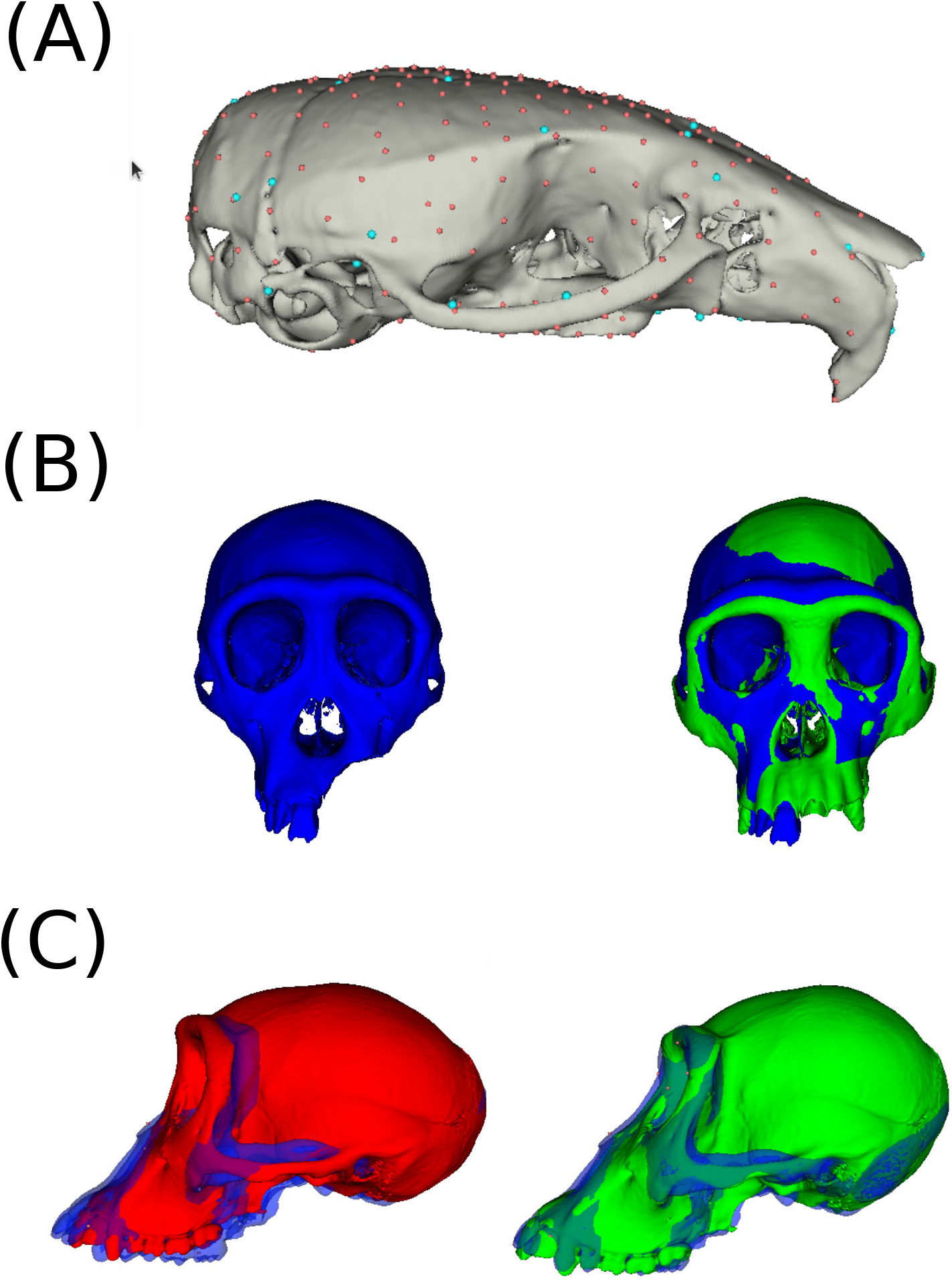
Non-standard uses of the ALPACA pipeline. (A) ALPACA can be used synergistically with other SlicerMorph modules to densely characterize the skull shape differences across individuals. In this case, a mouse skull mesh is illustrated with 51 type I landmarks (blue) plus 200 semi-landmarks automatically generated with PseudoLMGenerator from SlicerMorph. These landmark sets can be directly transferred to another specimen using ALPACA, therefore greatly reducing the landmarking effort. (B) When properly tuned, the ALPACA deformable registration step is able to deal (to some extent) with extra/missing parts. In this case, a Pan target mesh is missing a large part of the premaxilla and the maxilla. The warped source mesh is illustrated in green. Note that no significant artifact is present in the warped source mesh due to the missing craniofacial elements. Such scenario (missing elements) would be expected in a paleontological context. (C) Another possible use of ALPACA would be in an macroevolutionary context. In this case, we are illustrating the use of a source mesh from one genus (Pan; red) to predict landmark positions for another genus (Gorilla, blue). Note how the warped source mesh (green) has much clearer facial prognathism, matching the Gorilla mesh.

## Notes

https://github.com/SlicerMorph/SlicerMorph

### Competing Interest Statement

The authors have declared no competing interest.

